# Size-selective mortality fosters ontogenetic changes in collective risk-taking behaviour in zebrafish, *Danio rerio*

**DOI:** 10.1101/2021.09.13.460027

**Authors:** Tamal Roy, Robert Arlinghaus

## Abstract

Size-selective mortality is common in fish populations and can operate either in a positive size-selective fashion or be negatively size-selective. Through various mechanisms (like genetic correlations among behaviour and life-history traits or direct selection on behaviour co-varying with growth rate or size-at-maturation), both positive- and negative size-selection can result in evolutionary changes in behavioural traits. Theory suggests that size-selection alone favours boldness, but little experimental evidence exists about whether and to what extent size-selection can trigger its evolution. Here we investigated the impact of size-selective mortality on boldness across ontogeny using three experimental lines of zebrafish (*Danio rerio*) generated through positive (large-harvested), negative (small-harvested) and random (control line) size-selective mortality for five generations. We measured risk-taking during feeding (boldness) under simulated aerial predation threat and in presence of a live cichlid. We found that boldness decreased with ontogenetic age under aerial predation threat, and the small-harvested line was consistently bolder than controls. Collective personality emerged post larval stages among the selection lines. In presence of a cichlid, the large-harvested line was bolder at the highest risk of predation. The large-harvested line showed higher variability and plasticity in boldness across life stages and predation risks. Collectively, our results demonstrate that size-selective harvesting may evolutionarily alter risk-taking tendency. Size-selection alone favours boldness when selection acts on small fish. Selection typical of fisheries operating on large fish favours boldness at the highest risk of predation and increases behavioural variability and plasticity. There was no evidence for positive size-selection favouring evolution of shyness.

## Introduction

Intensive harvesting of wildlife can evolutionarily alter the behaviour of animal populations (Andersen et al. 2018; Leclerc et al. 2017), which can have repercussions for ecological processes, and ecosystem services, such as capture fisheries (Allendorf and Hard 2009; Jorgensen et al. 2007). Fisheries is an example of intensive harvesting which not only elevates adult mortality over natural levels, but is often selective of body-size and behavioural traits, like activity and boldness (Arlinghaus et al. 2017; Heino et al. 2015). While selection acting on large-sized fish (positive size-selection) is common in most commercial and recreational fisheries due to the selective properties of the gear or in response to harvest regulations (Heino et al. 2015; Monk et al. 2021), small-sized fish may also be preferentially caught due to regulations limiting the body-size in the landings (Ahrens et al. 2020) or be predated upon selectively by natural predators (negative size-selection) (Sogard 1997; Urban 2007). Both positive and negative size-selective mortality induced by fisheries or through natural predation may lead to the evolution of altered life-history and may also alter behavioural and physiological traits in exploited fish populations (Hollins et al. 2018; Matsumura et al. 2011; Renneville et al. 2020; Uusi-Heikkilä et al. 2008). While empirical evidence exists for the evolution of a fast life-history due to intensive size-selective mortality (Jorgensen et al. 2007; Uusi□Heikkilä et al. 2015; van Wijk et al. 2013), the evolutionary direction in which behavioural traits like boldness change in response to either elevated and unselective, or elevated and size-selective mortality has so far been studied only in a few models (Andersen et al. 2018; Claireaux et al. 2018; Jørgensen and Holt 2013). Animal personality expression typically changes through development (Bengston and Jandt 2014; Cabrera et al. 2021) and the question of how elevated and size-selective mortality may affect change in behaviour through ontogeny has no empirical answer. In this study, we aim to investigate the impact of size-selective mortality on boldness through ontogeny using experimental evolution in zebrafish (*Danio rerio*), a laboratory model.

Intensive fishing as well as natural mortality is typically size-selective in nature, meaning that fish of certain size-classes are preferentially harvested from populations (Hamilton et al. 2007; Heino et al. 2015; Jørgensen and Holt 2013). Most fishing gears preferentially capture the large and fast growing fish in populations (Law 2007), while most natural predators target smaller size classes (Stige et al. 2019; Urban 2007). For example, larger fish are typically harvested by trawls (Diaz Pauli et al. 2015; Kuparinen et al. 2009) or hook-and-line (Lewin et al. 2006). Elevated mortality of adults alone, even if unselective for size, is known to generally select for a fast life-history characterized by rapid maturation, rapid juvenile growth, elevated reproductive investment, reduced adult growth and reduced longevity (Hamilton et al. 2007; Heino et al. 2015; Jørgensen and Holt 2013; Laugen et al. 2014; Uusi□Heikkilä et al. 2015). Positive size-selection reinforces such life-history adaptations and adds pressures to mature early and at small sizes at the expense of post-maturation growth rate (Andersen et al. 2018; Jorgensen et al. 2007). Following the pace-of-life hypothesis, life-history adaptations towards a fast life-history should be related with increased boldness (Biro and Stamps 2008; Réale et al. 2010) to accumulate resources for rapid growth and fast sexual maturation (Jørgensen and Holt 2013; Montiglio et al. 2018). Indeed, population models that integrate behavioural processes have suggested that unselective (Claireaux et al. 2018; Jørgensen and Holt 2013) and size-selective harvesting would bring about increased boldness in exploited fish across a large gradient of size-selectivity (Andersen et al. 2018). However, Andersen et al. (2018) have also predicted that if size-selection is directed towards adult fish much larger than size-at-maturation, then this could lead to the evolution of shy behavioural phenotypes even under purely size-selected fisheries without any additional direct selection on behaviour. Since fishing gears are not just selective of body-size but also selective for behavioural traits such as boldness, trait-selective harvesting can favour shy behavioural phenotypes (timidity syndrome: Arlinghaus et al. 2017). While theory predicts the evolution of both elevated or reduced boldness with the evolution of fast life-history (Andersen et al. 2018), there is no experimental evidence to support either. Laboratory (Polverino et al. 2018) and field studies (Dhellemmes et al. 2021) have shown that a positive association between boldness and life-history based on POLS hypothesis (Réale et al. 2010) might break if populations are exposed to high adult mortality. Thus, divergent selection for life-history and behavioural traits are possible (Laskowski et al. 2021), and a positive correlation between life-history and boldness in fish populations adapted to positive size-selective harvesting may no longer be detectable.

A fast life-history evolves to cope with high adult mortality, thereby allowing fish reproducing early in life. Fish genetically predisposed to exert a fast-life history are expected to be bolder as juveniles because they must acquire the resources necessary for development of gonads early in life and therefore take risks during foraging (Claireaux et al. 2018; Jørgensen and Holt 2013). But fish with fast life-history are smaller post-maturation as a trade-off with elevated reproductive investment compared to the fish with slow life-history (Dunlop et al. 2009; Uusi□Heikkilä et al. 2015). Natural mortality due to predation is much higher for small sized juvenile fish than for large sized adults (Gislason et al. 2010; Lorenzen 2000). Thus, fish genetically programmed to exert a fast-life history may be expected to be more risk-averse as juveniles than as adults (Ballew et al. 2017). Hence boldness may change with transition through life-history stages among size-selected fish. Indeed, animal personality traits are associated with maturation and tend to be stable within, but not across life stages (Cabrera et al. 2021; Groothuis and Trillmich 2011). Studies across different fish species have revealed change in personality across developmental stages. For instance in mosquitofish *Gambusia holbrooki*, no evidence of personality was found in juveniles but personality emerged in subadult stage (Polverino et al. 2016a). Consistent differences in boldness and exploration increased during early ontogeny and reached an asymptote near sexual maturity in mangrove killifish *Kryptolebias marmoratus* (Edenbrow and Croft 2011). Behaviour tends to be repeatable within life stages. For example in tropical reef fish *Pomacentrus amboinensis*, juveniles immediately after transition from the larval stage were consistently bold across different time scales and contexts (White et al. 2015). In wild zebrafish, risk-taking tendency to feed was found to be consistent in adults over time (Roy and Bhat 2018) and across contexts (Roy et al. 2017). What is less known is whether exposure to multiple generations of intensive size-selection alters ontogenetic trajectories of fish behaviour. Previous studies have shown that size-selective mortality evolutionarily alters group risk-taking (Sbragaglia et al. 2020) and schooling (Sbragaglia et al. 2021) behaviour in adult zebrafish with experiments conducted at two time points, and only a recent study investigated the impact of size-selective mortality on change in shoaling behaviour through ontogeny (Sbragaglia et al. 2021, *unpublished*). Here we investigate if size-selective mortality affects change in group risk-taking behaviour across life-history stages in zebrafish. We measure group rather than individual phenotypes because zebrafish is a gregarious species (Suriyampola et al. 2016) and considering group behaviours are more ecologically relevant than studying zebrafish in isolation.

We use three experimental lines of zebrafish generated through positive (large-harvested), negative (small-harvested) and random (control) size-selective mortality for five successive generations to investigate the impact of size-selection on boldness through ontogeny. The experimental lines differ in life-history (Uusi□Heikkilä et al. 2015) and behavioural traits (Roy et al. 2021; Sbragaglia et al. 2019a; Sbragaglia et al. 2021; Sbragaglia et al. 2020; Uusi□Heikkilä et al. 2015) and these adaptations have genetic underpinnings, i.e., are truly genetic and not only adaptations within the realm of phenotypic plasticity (Sbragaglia et al. 2020; Uusi□Heikkilä et al. 2017; Uusi□Heikkilä et al. 2015). The large-harvested line mimics populations in most exploited fisheries where large-sized individuals are predominantly harvested. This line evolved a fast life-history characterized by smaller terminal body size and weight, early maturation and higher relative fecundity relative to the control line fish (Uusi□Heikkilä et al. 2015). The small-harvested line resembles fisheries where maximum-size limits exist or natural predation where mainly the smallest size classes are eaten. This line demonstrated reduced reproductive allocation but reached a similar terminal body-size compared to the control line, suggesting evolution towards a slow life-history (Uusi□Heikkilä et al. 2015). Both selection lines evolved altered maturation schedules by maturing at smaller size and younger age compared to the control line fish (Uusi□Heikkilä et al. 2015). Large-harvested line showed increased variation in body-size under different feeding environments (Uusi-Heikkilä et al. 2016). With respect to behaviour, juveniles (30 day old) of the small-harvested line were more explorative and bolder in an open field test than juveniles of large-harvested and control lines (Uusi□Heikkilä et al. 2015). Adult females of the small-harvested line showed decreased activity and boldness in an open field test and were less sociable than controls while large-harvested line females did not show differences with respect to the controls (Sbragaglia et al. 2019a). But there is critique of the open field test as a measure of boldness, and alternative tests have shown different results (Perals et al. 2017; Yuen et al. 2017). Groups of adults (230 and 240 day old) of the small-harvested line were found to take significantly more risk to feed under simulated predation threat while groups of the large-harvested line did again not differ in boldness from the control line (Sbragaglia et al. 2020). Adults (150 and 190 day old) of the large- and small-harvested line also formed more and less cohesive shoals compared to the control line fish (Sbragaglia et al. 2021). These studies were focused on one development stage and did not look at ontogenetic development. Here we measured group risk-taking behaviour (boldness) in the zebrafish lines in two different experiments across development to provide a complete picture on the evolution of boldness in response to size-selection.

Group risk-taking behaviour is a collective personality trait, and a shift in collective behaviour is expected through development since the needs of the group change with maturation (Bengston and Jandt 2014). We first investigated how collective risk-taking to feed (boldness) under simulated aerial predation threat (Sbragaglia et al. 2020; Ward et al. 2004) changed through ontogeny (i.e. from larval to adult stages) by measuring cumulative time spent by groups of fish feeding on the surface (defined as a risk zone) after releasing a model of a bird overhead (Sbragaglia et al. 2020). Fish may respond differently to aerial and aquatic predators as shown in different species because different species of predators exert conflicting selection pressures (Godin and Clark 1997; Templeton and Shriner 2004; Wund et al. 2015). Exposing fish to predation threat is important to unravel boldness and acquire reliable results (Klefoth et al. 2012). Therefore, we also tested boldness in presence of a predatory fish, a convict cichlid (*Amatitlania nirgrofasciata*) (Sailer et al. 2012; Toms and Echevarria 2014). We employed a contextual reaction norm approach (Dingemanse et al. 2010; Stamps and Groothuis 2010) by testing zebrafish in three contexts that differed in the kind of risk cues (visual and/or olfactory) perceived by the focal fish from the cichlid, and in a control context without the cichlid. Previous studies have explored behavioural reaction norms among fish by measuring behaviour across different time points or contexts (Killen et al. 2016). For example, three-spined sticklebacks (*Gasterosteus aculeatus*) showed consistent tendency to be bold or shy over a period of five weeks, and were less active and shyer when predation risk was higher (Fürtbauer et al. 2015). Wild zebrafish from high-predation habitats were consistently bold while fish from low-predation habitats were consistently shy across four contexts that needed fish to feed under varied risks of predation (Roy et al. 2017). In this study we measured the behaviour of groups of zebrafish across four contexts that differed in the risk of predation, and across ages >3 and >4 month.

Given that the large-harvested line is of a fast life-history and the small-harvested line is of a slow fast-life history (Uusi□Heikkilä et al. 2015), and considering that Andersen et al. (2018) showed that size-selection tends to generate bold fish unless only the largest fish are selected, we hypothesized that both positive and negative size-selection would lead to elevated boldness compared to the controls and that this effect would be more pronounced for the negatively size-selected line. We would expect that maturation will change boldness, with fish among the large-harvested line being particularly bold in the juvenile stage and be less bold after transition to the mature stage to adjust behavior to the now smaller body-size. We further expected that the two size-selected lines would adjust their boldness behavior in response to the degree of apparent predation risk thereby showing different levels of behavioural variability and plasticity. Disruptive selection due to harvesting may alter trait variability (Landi et al. 2015). For example in pike (*Esox lucius*), fishery increases variability in somatic growth rate and size at age (Edeline et al. 2009) while positive size-selective mortality leads to increased variation in body-size in experimentally harvested populations of zebrafish (Uusi-Heikkilä et al. 2016). Here we expect that the plastic behavioural adaptation would be more pronounced in the large-harvested line because of their internal conflict between reaping resources through foraging and avoiding being predated upon due to their smaller body-size. By contrast, we expected the small-harvested line to be consistently bolder than controls and show less behavioural plasticity against the gradient of predation risk as this line has been found to be bold (Sbragaglia et al. 2020) and is larger in size meaning that the predation risk is generally lower.

## Material and Methods

### Selection lines

We used F_16_ of the selection lines of zebrafish (large-, random- and small-harvested lines, each with a replicate, i.e. six lines in total), described in (Uusi□Heikkilä et al. 2015). These lines were produced by subjecting a wild-population of zebrafish to intensive selection (i.e. 75% harvest rate per generation) for five successive generations (F1 to F5), and thereafter stopping selection for the subsequent generations to remove maternal effects (Moore et al. 2019). 25% of the smallest and largest individuals were used as parents in successive generations in the large- and small-harvested line. Simultaneously, 25% of random individuals were selected for reproduction every generation to produce the control group. Fish were harvested in the subsequent generations based on when 50% of the control line fish became mature. The selection lines differed in body-size and life-history (Uusi□Heikkilä et al. 2015) and the evolved differences in body-size were maintained in F13 (Roy et al. 2021; Sbragaglia et al. 2019b) and F16 (ESM, Fig. S1). The selection lines also differed in broad scale gene expression (Sbragaglia et al. 2020; Uusi□Heikkilä et al. 2017) thereby confirming that the phenotypic differences had genetic underpinnings (Sbragaglia et al. 2020; Uusi□Heikkilä et al. 2015).

We housed the F_15_ fish of selection lines in laboratory in six round holding tanks (diameter: 79 cm, height: 135 cm, volume: 320l) at a density of approximately 1000 per tank under 12:12 Light: Dark cycle. The water temperature was maintained at 27°C by a circulation system, and fish were fed twice daily with commercial flake food (TetraMin Tropical). For producing the F16 fish, we stocked six 2 l spawning boxes (total 36) each with four males and two females (Roy et al. 2021; Uusi□Heikkilä et al. 2010) selected randomly from the F15 population, and allowed them to breed. We pooled the embryos produced from each line and transferred 80 embryos into 10 1.5 l boxes (eight per box). We used 480 fish (i.e. 8 × 60 groups; 10 groups per replicate line, 20 per treatment) in total and tracked the behaviour of fish through ontogeny. The fish were fed twice a day with powdered flake food in the larval and juvenile stages, and like F15 fish when adult.

### Risk-taking under simulated aerial predation

We tested risk-taking to feed under simulated aerial predation threat among groups of fish across selection lines using a tank diving paradigm employed previously by Sbragaglia et al. (2020). Collective risk-taking behaviour is a repeatable trait in zebrafish (Sbragaglia et al. 2019b; Sbragaglia et al. 2019c) and is a group level personality trait (Bengston and Jandt 2014; Jolles et al. 2018). We tested 60 groups (10 from each replicate line, 20 per treatment) or 480 fish in total. We conducted the assay every week from 8 to 22 dpf (larval stages), at 45 dpf (juvenile), 61 and 85 dpf (subadult stages), and then every three weeks till 148 dpf and after six weeks at 190 dpf (adult stages) (Alfonso et al. 2020). Behavioural changes in laboratory reared zebrafish is associated with morphological and physiological changes during larval (8 - 21 dpf), juvenile (21 – 60 dpf including metamorphosis around 45 dpf), subadult (60 – 90 dpf) and adult (90 – 190 dpf) stages (Alfonso et al. 2020; Stednitz and Washbourne 2020). Though the maturation schedules of our size-selected lines are different (Uusi□Heikkilä et al. 2015), we tested them at the abovementioned stages to have an uniform experimental timeline (Fig. 1).

**Fig. 1:**
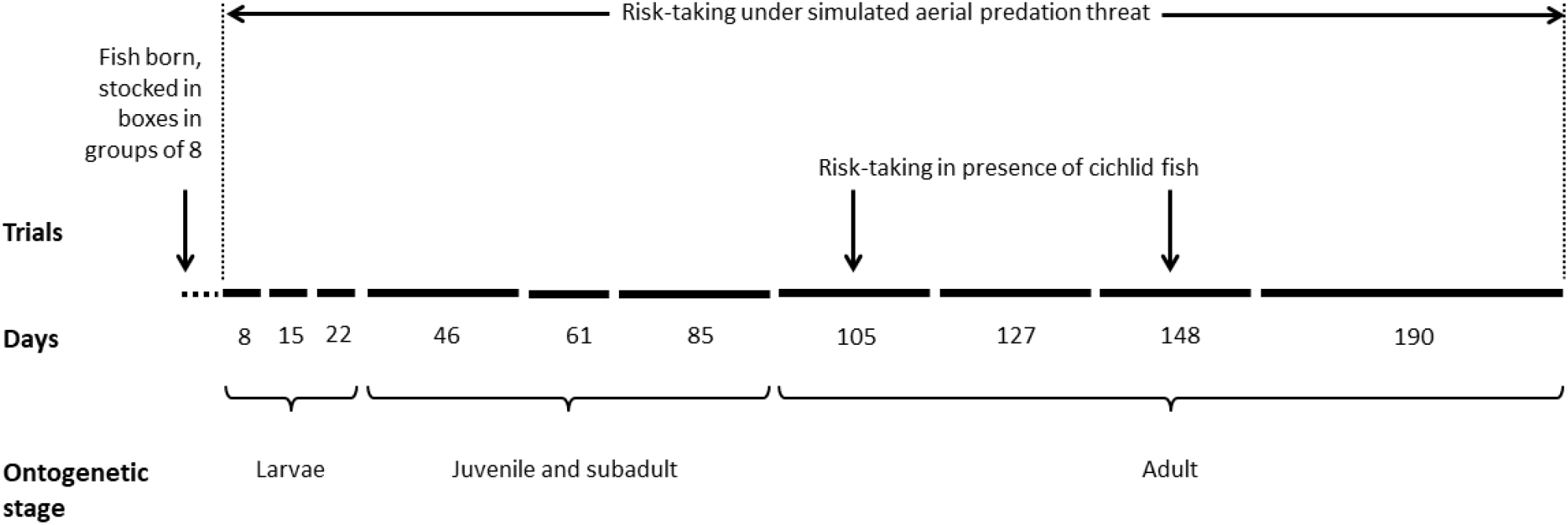
Experimental timeline. Fish across selection lines were tested for risk-taking behaviour in presence of an aerial predation threat from age 8 to 190 dpf, and in presence of a cichlid fish at > 3- and > 4-month age.

We used a rectangular glass tank (30 × 10 × 25 cm) as the experimental setup with white opaque walls on three sides and placed the setup behind a white curtain to avoid external disturbances affecting fish behaviour (Fig. 2a). The tank was filled with system water up to a level of 20 cm and we demarcated 4 cm from the top as the ‘surface zone’. We gently transferred a group of eight fish from their holding into the arena and started recording their behaviour with a webcam (LogitechB910) installed 20 cm from the transparent side of the setup. After 2 min, we added food to the surface of water and allowed the fish to feed for 30 sec. We released a paper cutout of bird (simulated predator) at a height of 10 cm from the water surface for 15 sec (Fig. 2a) and after retrieving it back, we allowed the fish to resume feeding for 5 min. From the video recordings, we scored the cumulative time spent by fish at the surface (Egan et al. 2009; Kalue 2017) while feeding after we retrieved the predator model as a measure of boldness.

**Fig. 2:**
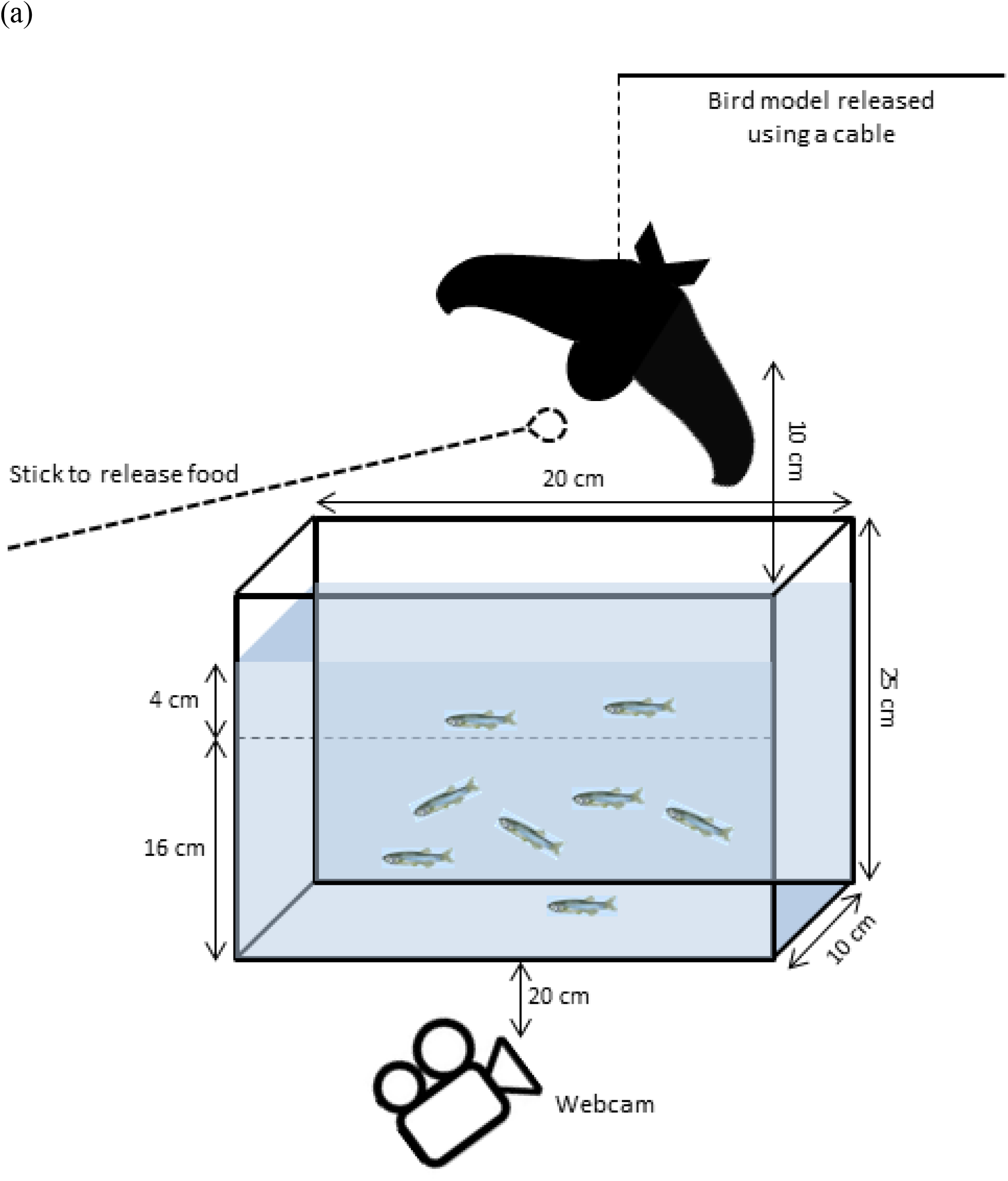
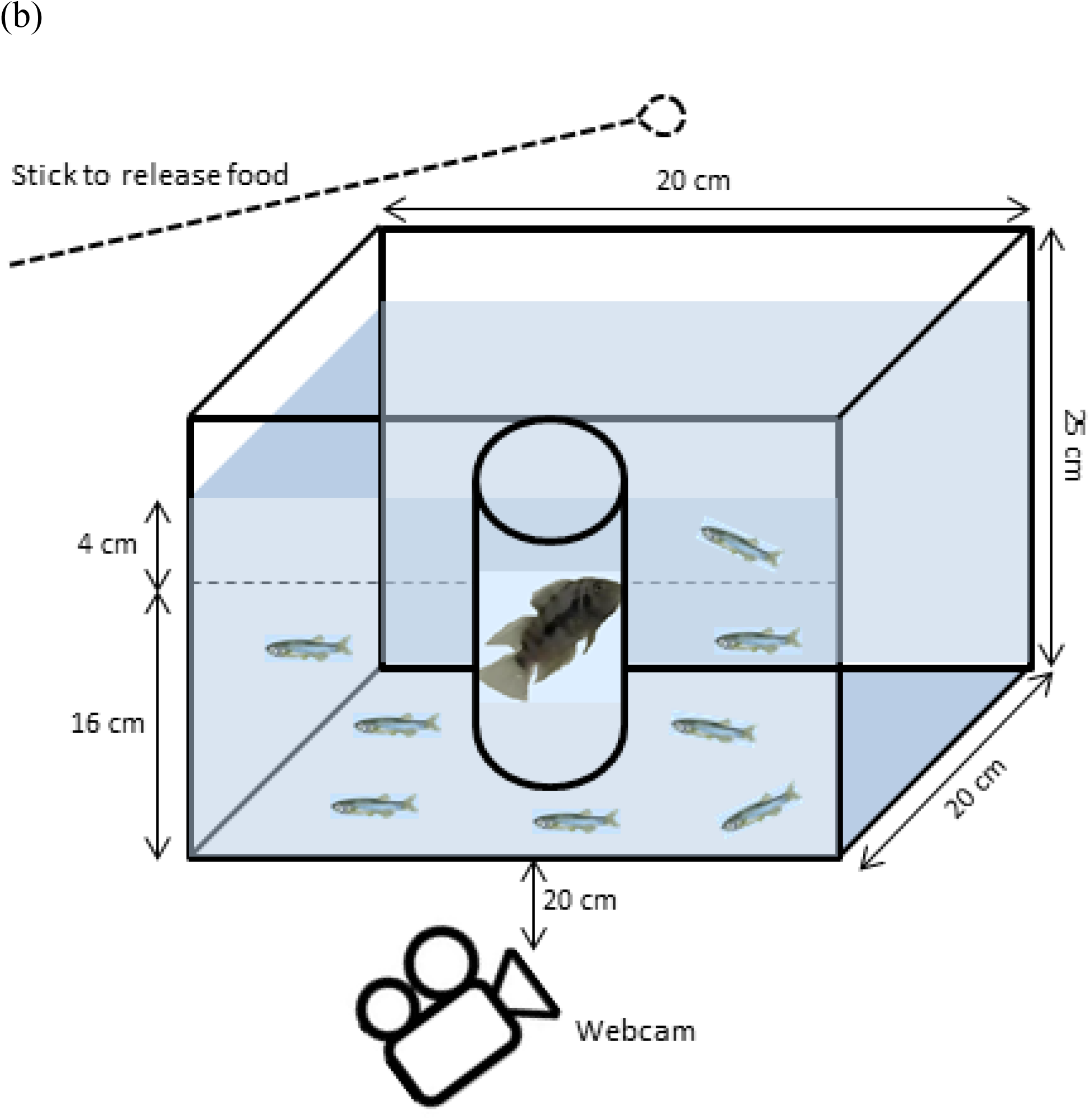
Experimental setups for testing risk-taking to feed in presence of (a) simulated aerial predator (in the form of a paper cutout of bird), and (b) a live convict cichlid fish. The image of zebrafish have been taken from Guerreiro (2008).

### Risk-taking in presence of a live predator

We tested contextual norms (Stamps and Groothuis 2010) for risk-taking to feed in presence of an aquatic predator, a convict cichlid (*Amatitlania nirgrofasciata*), twice (at > 3 and > 4 month age) among adult zebrafish groups. Previous studies testing risk-taking behaviour in zebrafish have used convict cichlids (Sailer et al. 2012; Toms and Echevarria 2014). We tested 42 groups (7 from each replicate line, 14 per treatment) or 336 fish in total. We used a similar setup like previous with a rectangular glass tank (30 × 20 × 25 cm) having three opaque walls (Fig. 2b) and a demarcated surface zone. We tested fish across three contexts (contextual norms) where the zebrafish could only see, only smell, and both see and smell the live predator. For this, we introduced a cichlid fish into a cylindrical container that was transparent (for visual cue) and permeable (for both visual and chemical cues) or opaque and permeable (for chemical cue), and placed the container in the experimental arena (Fig. 2b). We also tested fish in a controlled setting without the predator. During the experiment, we transferred a group of eight fish into the arena and started recording their behaviour. After 2 min, we added food on the water surface and allowed the fish to feed for 5 min. From the video recordings, we scored the cumulative time spent by zebrafish at the surface.

## Statistical analysis

We constructed linear mixed-effects regression models (lmer) to test for difference in boldness among selection lines through ontogeny under aerial predation threat, and across different degrees of risk (contexts) in presence of live predator. In tests for risk-taking under simulated aerial predation, we first transformed the response variable (cumulative time spent at the surface) using cube-root transformation. We fitted mixed effects models using the transformed measure as a dependent variable, interaction of ‘Selection line’ (large-harvested, control and small-harvested) and ‘Age’ (ontogenetic stage) as the fixed effect and ‘Group ID’ nested within ‘Replicate’ (two per line) as the random effect. To test for consistency in boldness across life-history stages, we estimated the adjusted repeatability (Nakagawa and Schielzeth 2010) for each selection line over three stages; larval (8 to 22 dpf), juvenile and subadult (46 to 85 dpf) and adult (105 to 190 dpf) using the ‘rpt’ function. We considered the cube-root transformed measure as dependent variable, ‘Age’ as fixed effect, and ‘Group ID’ as random effect. We used 95% confidence intervals with a significance level of 5% as estimates of uncertainty. To estimate variability and plasticity in boldness in each line, we used between-group and within-group variances (Polverino et al. 2016a; Roy et al. 2017) obtained by running separate mixed-effects regression models at each stage.

To investigate differences in boldness across contexts in presence of a live predator, we first transformed the response variable (cumulative time spent at the surface) using Tukey’s Ladders of Powers transformation. We then fitted mixed effects models using the Tukey-transformed measure as the dependent variable, interaction of ‘Selection line’ and ‘Context’ (control, visual, chemical and visual+chemical) as the fixed effect, ‘Group ID’ nested within ‘Replicate’ as the random intercept and ‘Age’ as the random slope. To test for consistency in boldness across four contexts, we estimated the adjusted repeatability (with 95% CI) like previous for each selection line by considering the Tukey-transformed measure as dependent variable, interaction of ‘Context’ and ‘Age’ as fixed effect, and ‘Group ID’ as random effect. To estimate behavioural variability and plasticity in each line, we similarly obtained between- and within-group variances by running separate mixed-effects regression models.

All analyses were conducted in R version 3.6.1 (R Development Core Team 2019). Data were transformed using the ‘*rcompanion*’ package (Mangiafico and Mangiafico 2017), mixed effects models were constructed using ‘*lmerTest*’ package (Kuznetsova et al. 2017) and behavioural repeatability was estimated using ‘*rptR*’ package (Stoffel et al. 2017). Box-whisker plots were made using ‘*ggplot2*’ (Wickham 2011) and ‘*ggpubr*’ (Kassambara and Kassambara 2020) packages in R.

## Results

We found a significant and line-specific ontogenetic effect on boldness (selection line × age; F18, 513 = 5.32, p<0.01) (Table 1a) meaning that boldness (risk-taking behaviour) changed significantly through ontogeny across selection lines (Fig. 3a). The cumulative time spent by fish at the surface decreased significantly as the fish matured and developed through ontogeny, i.e., bigger and older fish generally took less risk to feed than smaller and younger individuals (Fig. 3a-d, 4). The small-harvested (SH) line fish were significantly bolder than the control (random-harvested) line fish at larval stage 22 dpf (t=2.21, p=0.03) (Fig. 3b), subadult stages 61 dpf (t=1.77, p=0.08) and 85 dpf (t=3.05, p<0.01) (Fig. 3c), and adults stages 105 dpf (t=4.08, p<0.01), 127 dpf (t=6.82, p<0.01), 148 dpf (t=2.47, p=0.01) and 190 dpf (t=3.06, p<0.01) (Fig. 3d). The large-harvested line fish did not differ in boldness from the control line fish at any stage (Fig. 3a-d, Table 1b). Hence, while all lines decreased their boldness levels as the fish aged (Fig. 3a), the decrease in boldness after maturation was less pronounced in the small-harvested line (Fig. 3a, d). Boldness levels decreased more in the control and large-harvested line fish and they became particularly shy after maturation compared to the juvenile and larval stages (Fig. 3a-d).

**Table 1:** Results of (a) main effects and (b) fixed effects terms obtained from linear mixed effects model comparing boldness in fish from selection lines LH (large-harvested) and SH (small-harvested) with the control line across ontogenic stages A (8 dpf) to J (190 dpf). Significance of the fixed effects and their interactions are in bold (marginal: ‘^+^’).

| (a) Main effects | Sum Sq | Mean Sq | NumDF | DenDF | F value | Pr(>F) |
| --- | --- | --- | --- | --- | --- | --- |
| Selection line | 7.56 | 3.780 | 2 | 3.00 | 4.65 | 0.12 |
| Age | 1399.40 | 155.49 | 9 | 513.16 | 191.42 | **< 0.001** |
| Selection line × Age | 77.73 | 4.32 | 18 | 513.16 | 5.317 | **< 0.001** |

| (b) Fixed effects | Estimate | Std. Error | df | t value | Pr(>\|t\|) |
| --- | --- | --- | --- | --- | --- |
| Intercept | 6.60 | 0.34 | 6.55 | 19.63 | **< 0.001** |
| Selection line: LH | -0.004 | 0.47 | 6.55 | -0.01 | 0.99 |
| Selection line: SH | 0.05 | 0.47 | 6.55 | 0.11 | 0.91 |
| Stage B (15 dpf) | -0.71 | 0.28 | 512.34 | -2.49 | **0.01** |
| Stage C (22 dpf) | -1.10 | 0.28 | 512.34 | -3.88 | **< 0.001** |
| Stage D (46 dpf) | -0.95 | 0.29 | 518.59 | -3.32 | **< 0.001** |
| Stage E (61 dpf) | -1.57 | 0.28 | 512.34 | -5.50 | **< 0.001** |
| Stage F(85 dpf) | -2.76 | 0.28 | 512.34 | -9.70 | **< 0.001** |
| Stage G (105 dpf) | -4.39 | 0.28 | 512.34 | -15.40 | **< 0.001** |
| Stage H (127 dpf) | -4.90 | 0.28 | 512.34 | -17.20 | **< 0.001** |
| Stage I (148 dpf) | -4.06 | 0.28 | 512.34 | -14.26 | **< 0.001** |
| Stage J (190 dpf) | -4.14 | 0.28 | 512.34 | -14.51 | **< 0.001** |
| LH × Stage B | 0.59 | 0.40 | 512.34 | 1.46 | 0.14 |
| SH × Stage B | 0.27 | 0.40 | 512.34 | 0.66 | 0.51 |
| LH × Stage C | 0.45 | 0.40 | 512.34 | 1.11 | 0.27 |
| SH × Stage C | 0.89 | 0.40 | 512.34 | 2.21 | **0.03** |
| LH × Stage D | -0.02 | 0.40 | 512.34 | -0.05 | 0.96 |
| SH × Stage D | 0.56 | 0.40 | 512.34 | 1.40 | 0.16 |
| LH × Stage E | -0.15 | 0.40 | 512.34 | -0.38 | 0.71 |
| SH × Stage E | 0.71 | 0.40 | 512.34 | 1.77 | 0.08<sup>+</sup> |
| LH × Stage F | 0.10 | 0.40 | 512.34 | 0.24 | 0.81 |
| SH × Stage F | 1.23 | 0.40 | 512.34 | 3.05 | **< 0.01** |
| LH × Stage G | -0.15 | 0.40 | 512.34 | -0.36 | 0.71 |
| SH × Stage G | 1.65 | 0.40 | 512.34 | 4.08 | **< 0.001** |
| LH × Stage H | 0.65 | 0.40 | 512.34 | 1.62 | 0.10 |
| SH × Stage H | 2.75 | 0.40 | 512.34 | 6.82 | **< 0.001** |
| LH × Stage I | -0.36 | 0.40 | 512.34 | -0.91 | 0.36 |
| SH × Stage I | 0.99 | 0.40 | 512.34 | 2.47 | **0.01** |
| LH × Stage J | 0.00 | 0.40 | 512.34 | 0.01 | 0.99 |
| SH × Stage J | 1.23 | 0.40 | 512.34 | 3.06 | **< 0.01** |

**Fig. 3:**
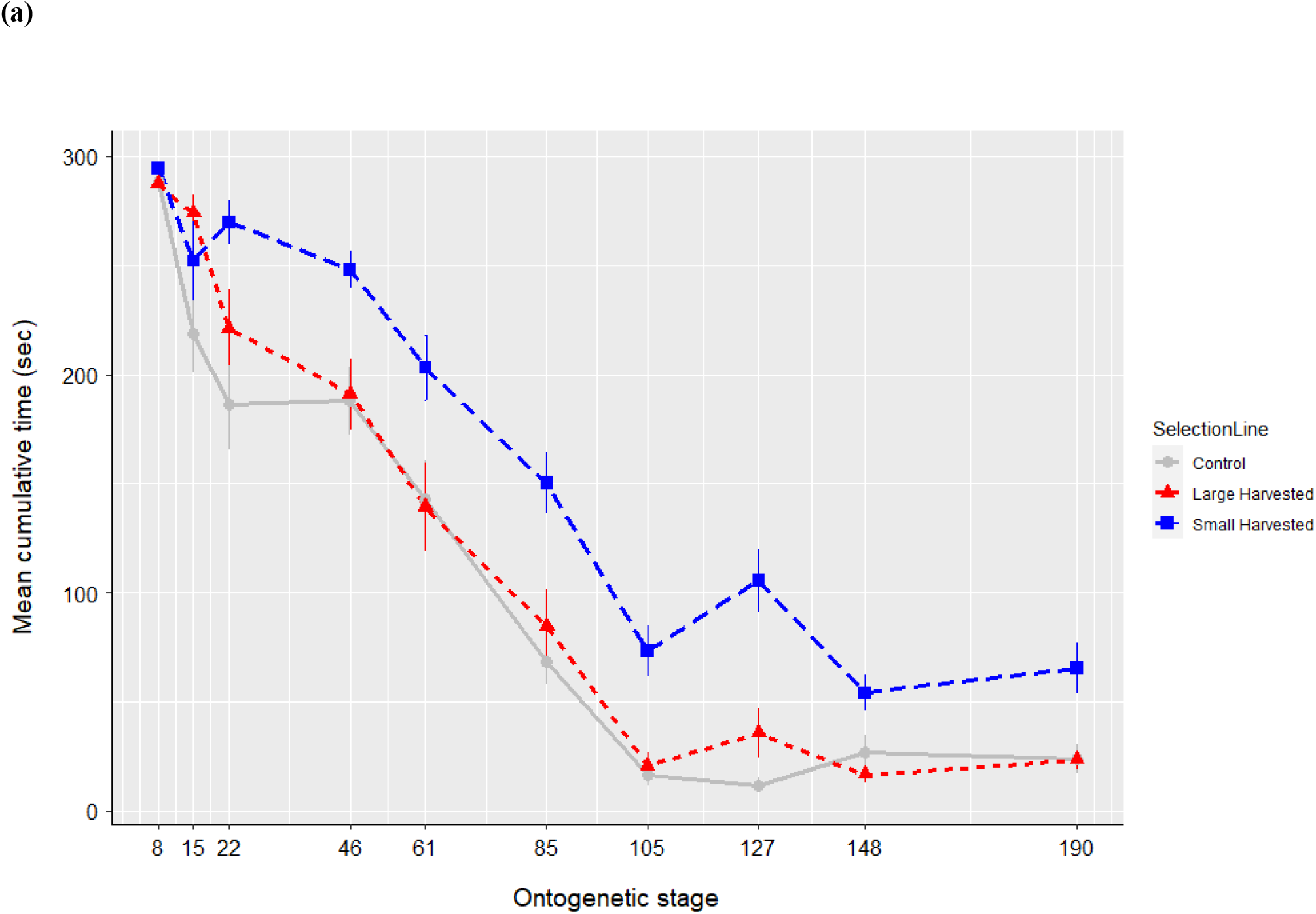
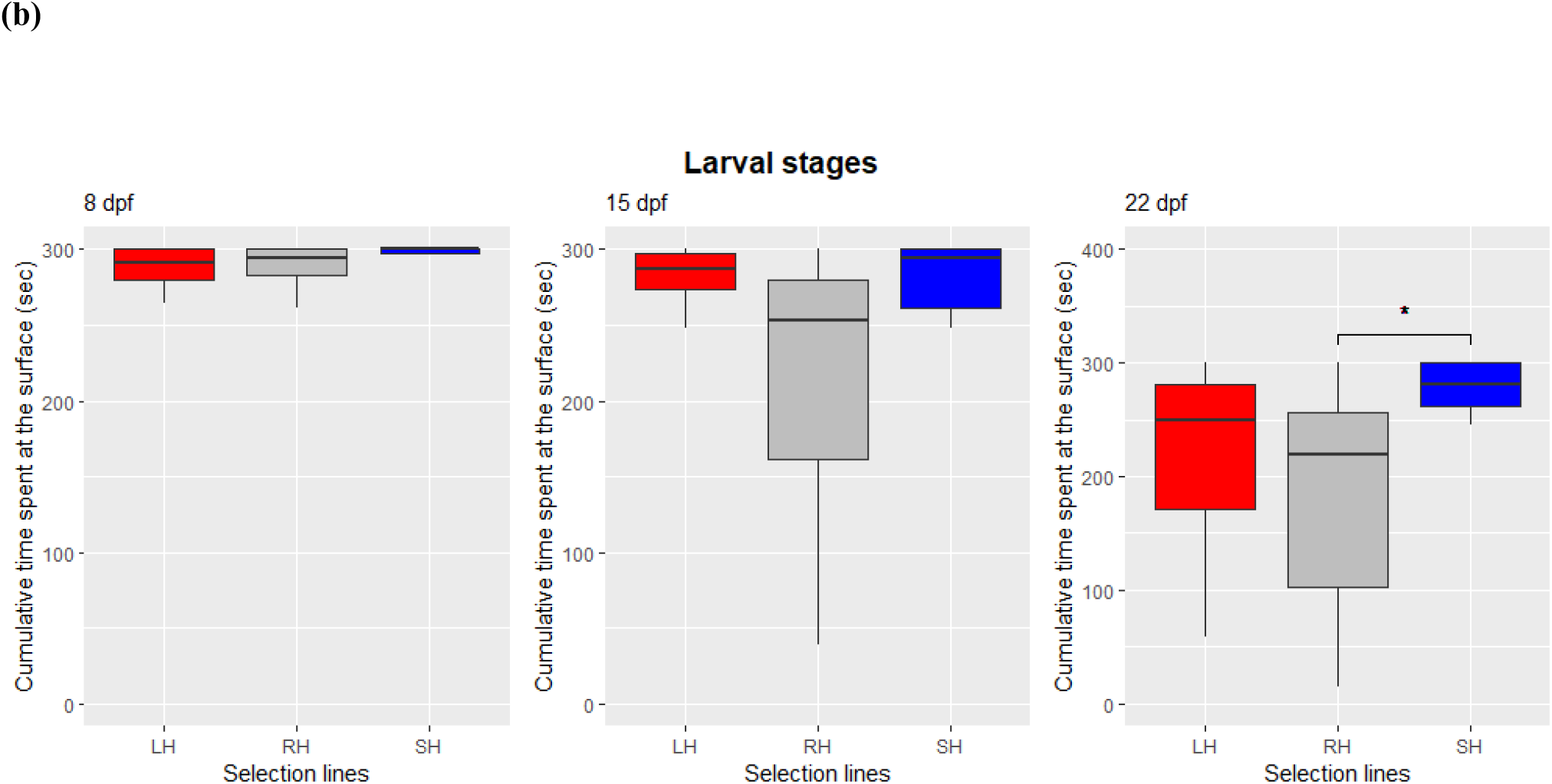
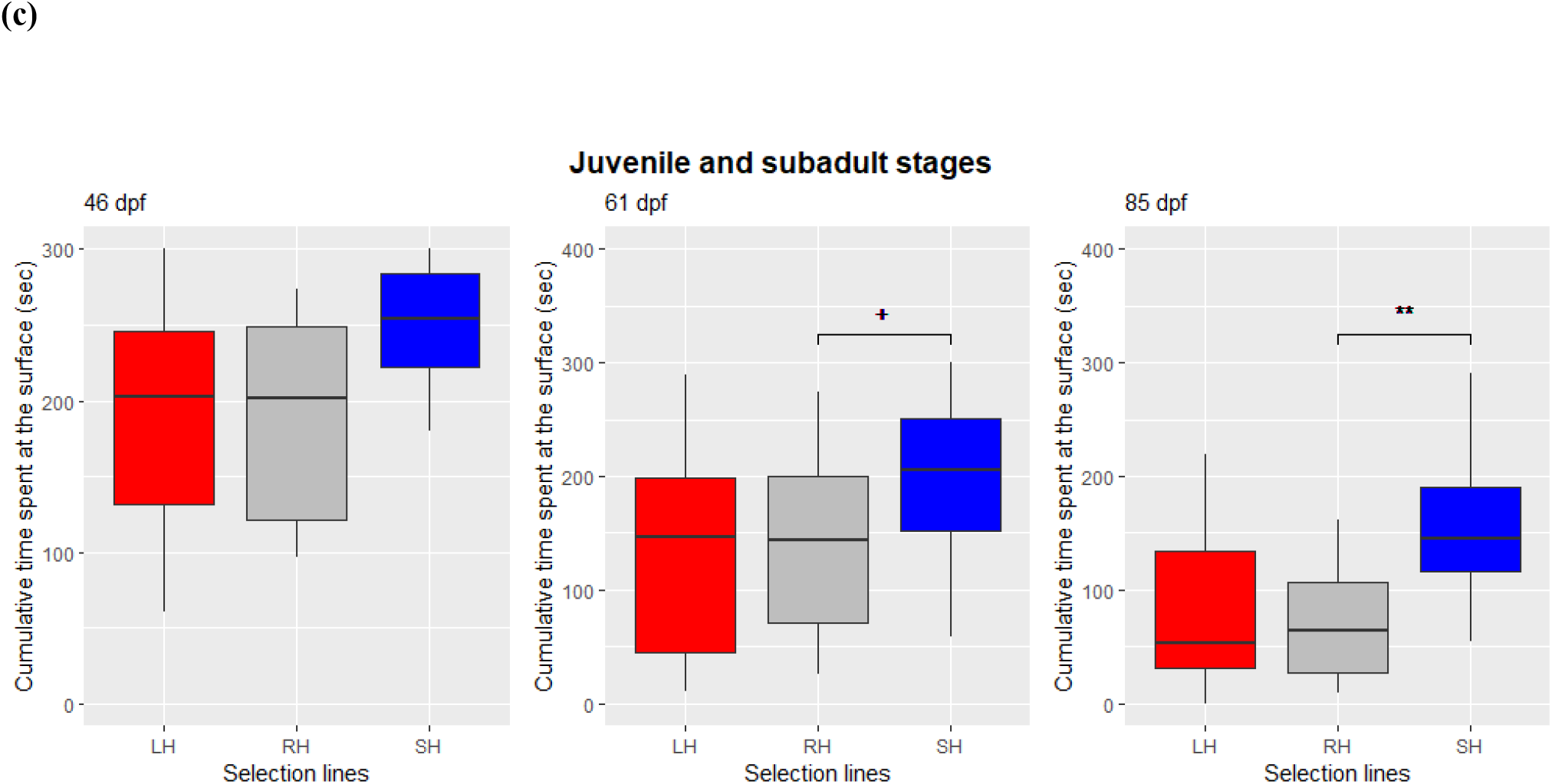
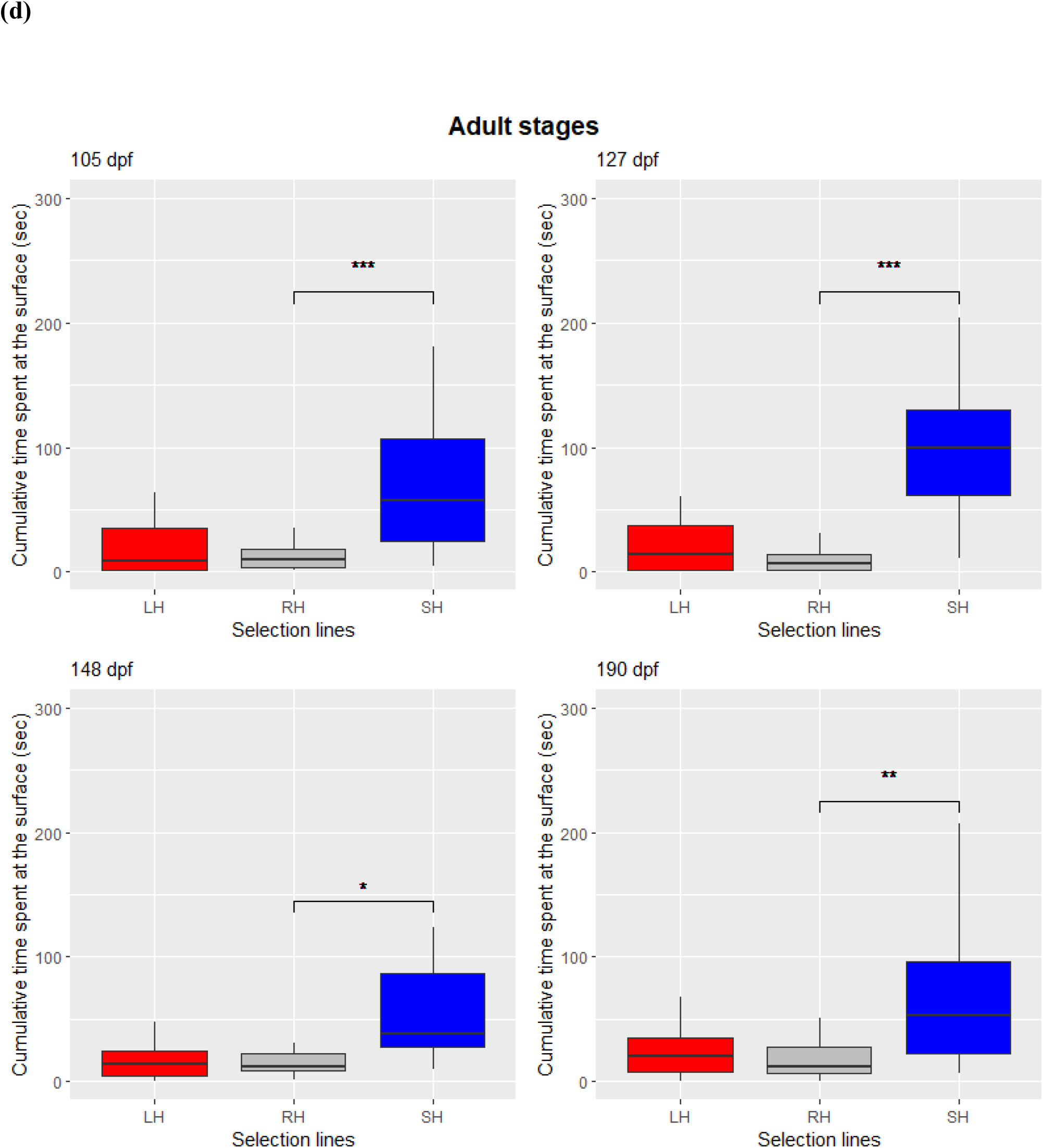
Change in boldness (measured as cumulative time spent at the surface) through ontogeny among large-harvested (LH: red), control (RH: grey) and small-harvested (SH: blue) selection lines. The first panel (a) shows change in mean cumulative time (±SE) spent at the surface across all ontogenetic stages (8 to 190 dpf). The second panel (b) shows behavioural change across larval stages from 8 to 22 dpf. The third panel (c) shows behavioural change across juvenile (∼ 46 dpf when metamorphosis is complete) and subadult stages at 61 and 85 dpf. The fourth panel (d) shows behavioural change across adult stages from 105 to 190 dpf. Significant differences are indicated with bars and codes *** (p<0.001), ** (p<0.01), * (p<0.05), and ^**+**^ (p<0.1).

**Fig. 4:**
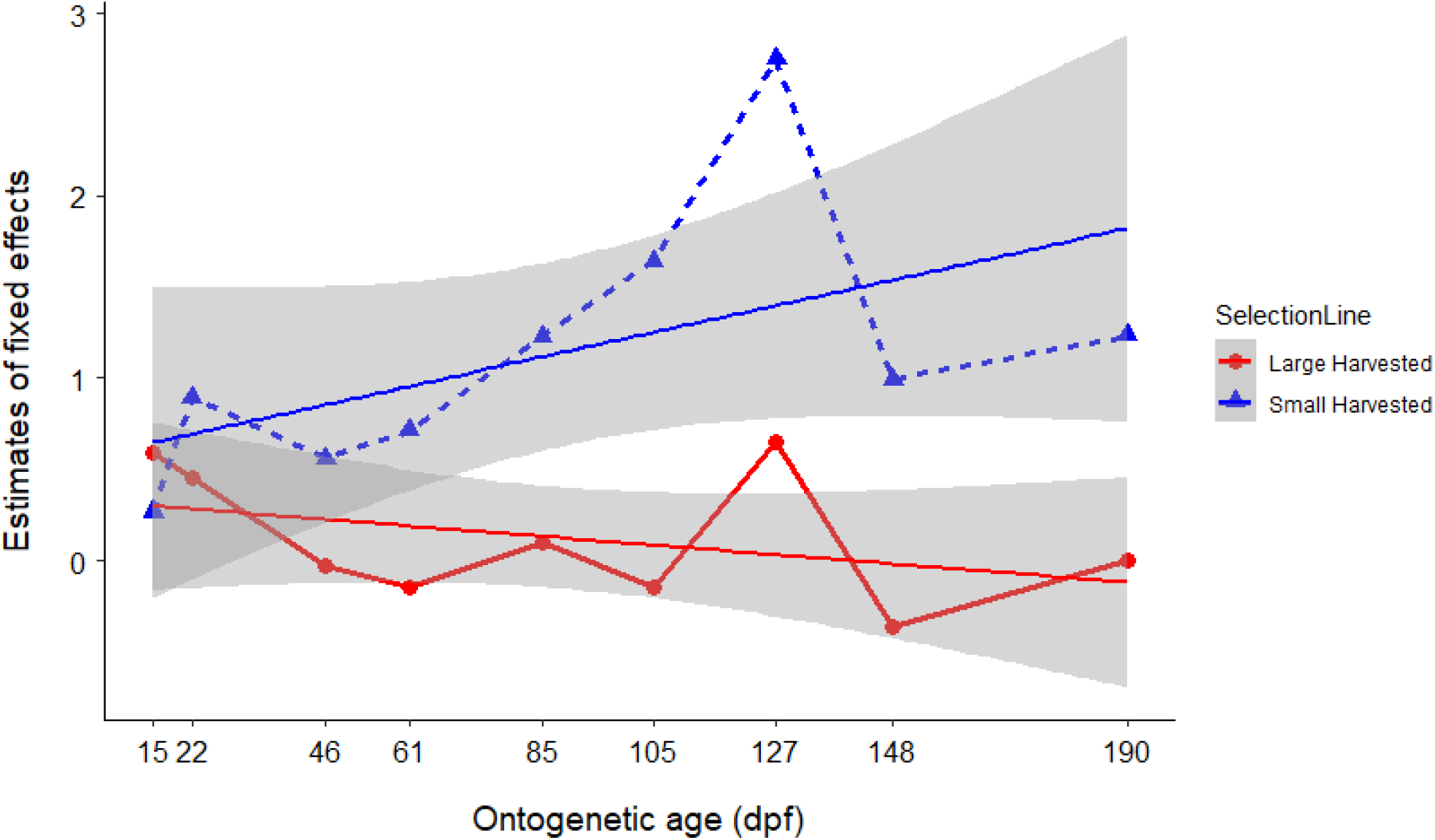
Estimates of the interaction of fixed effects ‘selection line’ (LH and SH) and ‘ontogenetic age’ (15 to 190 dpf) obtained from the results of linear mixed effects model.

In tests for behavioural consistency among life-history stages, we found that the selection lines showed no significant repeatability in boldness across larval stages i.e. from 8 to 22 dpf (Table 2a). All lines showed low between-group and within-group variances indicating low behavioural variability and plasticity among groups (Table 2a). Significant repeatability was observed in large-harvested (R=0.48, p<0.001) and small-harvested (R=0.43, p<0.001) lines while the control line fish showed no significant repeatability in boldness from juvenile (46 dpf) to subadult (85 dpf) stages (Table 2b). Large-harvested line showed higher between-group variance while small-harvested line showed lower between between-group variance compared to the controls (Table 2b). Large-harvested and control line fish showed high within-group variances i.e. high behavioural plasticity compared to the small-harvested line fish (Table 2b). In adults from 105 to 190 dpf, significant repeatability in boldness was observed in large-harvested (R=0.61, p<0.001), control (R=0.48, p<0.001) and small-harvested (R=0.57, p<0.001) lines (Table 2c). The large-harvested line showed high between-group and within-group variances indicating high behavioural variability and plasticity compared to small-harvested and control line fish (Table 2c).

**Table 2:** Between- and within-group variances and repeatability in boldness among selection lines across (a) larval (8 to 22 dpf), (b) juvenile and subadult (46 to 85 dpf), and (c) adult stages (105 to 190 dpf): estimates of repeatability with 95% confidence interval and significance level of 5%. Significant results are in bold.

| Selection line | Between-group variance | Within-group variance | R | SE | 95% CI | p |
| --- | --- | --- | --- | --- | --- | --- |
| (a) Larval stages |  |  |  |  |  |  |
| Large-harvested | 0.05 | 0.24 | 0.16 | 0.13 | 0, 0.45 | 0.12 |
| Random-harvested | 4.22e<sup>-16</sup> | 7.64e<sup>-1</sup> | 0.00 | 0.00 | 0, 0 | 0.5 |
| Small-harvested | 0.00 | 0.31 | 0.00 | 0.08 | 0, 0.28 | 1 |
| (b) Juvenile and sub-adult stages |  |  |  |  |  |  |
| Large-harvested | 0.77 | 0.83 | 0.48 | 0.14 | 0.19, 0.71 | **<0.001** |
| Random-harvested | 0.00 | 0.90 | <0.01 | 0.08 | 0, 0.28 | 1 |
| Small-harvested | 0.28 | 0.37 | 0.43 | 0.14 | 0.13, 0.66 | **<0.001** |
| (c) Adult stages |  |  |  |  |  |  |
| Large-harvested | 1.19 | 0.76 | 0.61 | 0.11 | 0.37, 0.78 | **<0.001** |
| Random-harvested | 0.55 | 0.59 | 0.48 | 0.12 | 0.24, 0.68 | **<0.001** |
| Small-harvested | 0.62 | 0.47 | 0.57 | 0.10 | 0.35, 0.75 | **<0.001** |

In the test for boldness in presence of a live predatory fish, we found an overall significant effect of context (F_3,243_ =2.64, p=0.05) (Table 2a) and a marginally significant interaction effect of LH line and ‘Visual & Olfactory’ context (t=1.88, p=0.06) (Table 2b). This meant that the large-harvested line fish spent more time at the surface and were bolder when the risk of predation was highest i.e. when zebrafish perceived both visual and olfactory cues from the cichlid (Fig. 5). Though not significant, the small-harvested line fish spent more time at the surface and were generally bolder across all contexts i.e. under all risks of predation compared to the controls (Fig. 5).

**Fig. 5:**
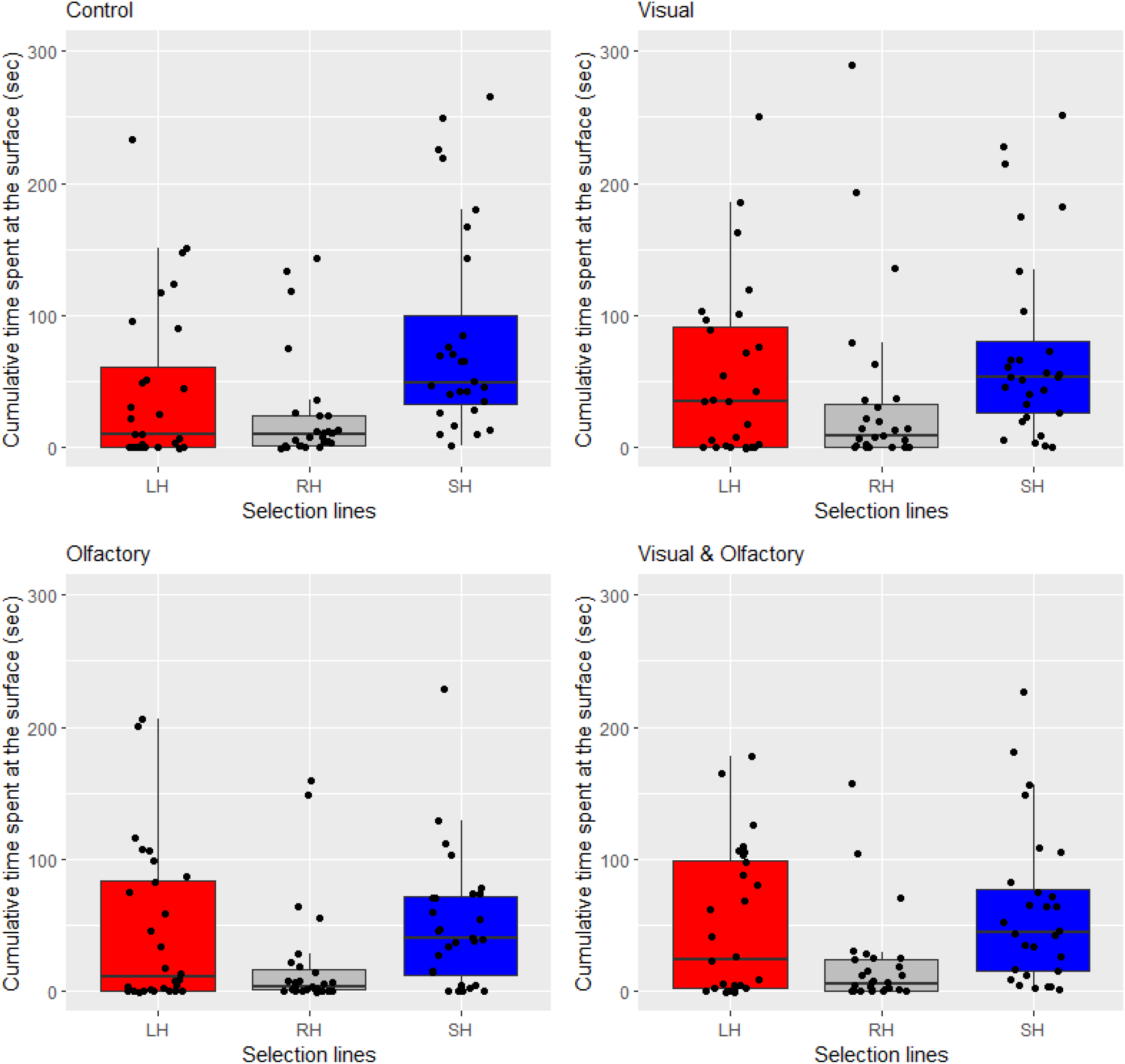
Combined plot (for > 3- and > 4-month age) of cumulative time spent at the surface (boldness) by fish among three selection lines (large-harvested, control and small-harvested) across three contexts where zebrafish perceived visual, olfactory and both visual and olfactory cues from the predator, and in a control setting without the predator.

In tests for behavioural consistency, we found significantly high repeatability in boldness across four contexts and across age among large-harvested (R=0.86, p<0.001), control (R=0.56, p<0.001) and small-harvested (R=0.54, p<0.001) line fish (Table 4). Large-harvested line fish showed high between-group variance indicating high behavioural variability and low within-group variance i.e. low plasticity, compared to control and small-harvested line fish (Table 4).

**Table 3:**
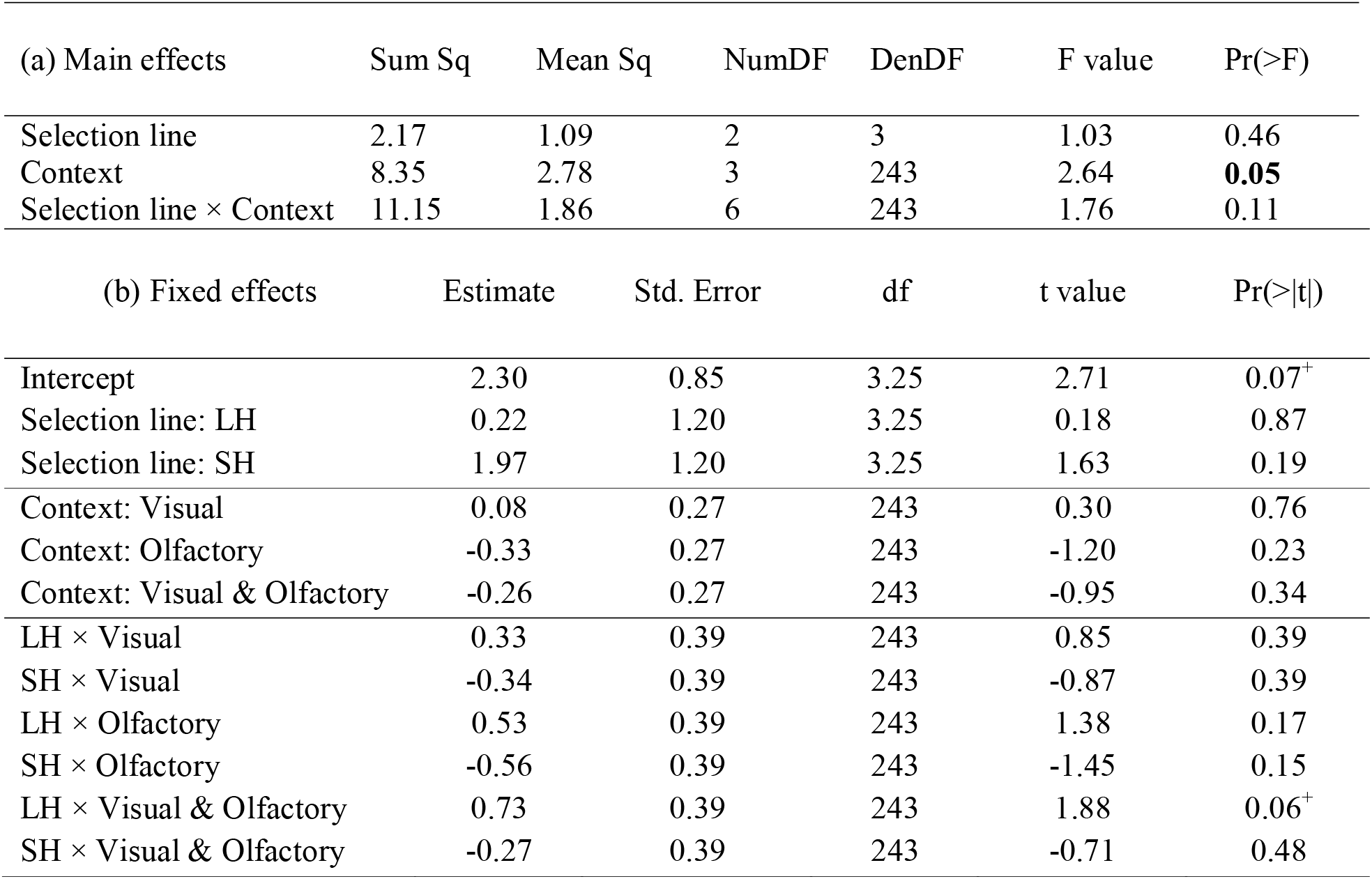
Results of (a) main effects and (b) fixed effects terms obtained from linear mixed effects model comparing boldness in fish from selection lines LH (large-harvested) and SH (small-harvested) with the control line across four contexts (‘Control’, ‘Visual’, ‘Olfactory’ and ‘Visual & Olfactory’). Significant result is in bold and marginal significance are indicated with ‘^+^’.

**Table 4:** Between- and within-group variances and repeatability in boldness across contexts among selection lines: estimates of repeatability are with 95% confidence interval and significance level of 5%. Significant results are in bold.

| Selection line | Between-group<br>variance | Within-group<br>variance | R | SE | 95% CI | p |
| --- | --- | --- | --- | --- | --- | --- |
| Large-harvested | 4.41 | 0.71 | 0.86 | 0.06 | 0.71, 0.93 | **<0.001** |
| Random-harvested | 1.71 | 1.34 | 0.56 | 0.11 | 0.28, 0.74 | **<0.001** |
| Small-harvested | 1.39 | 1.18 | 0.54 | 0.11 | 0.27, 0.73 | **<0.001** |

## Discussion

We found that five generations of size-selective harvesting left an evolutionary legacy on collective risk-taking behaviour (boldness) in zebrafish at different levels of predation risk. Boldness levels decreased with ontogenetic age under aerial predation threat and fish became shyer after maturation compared to the juvenile and larval stages. This effect was less pronounced in the small-harvested line compared to the control and large-harvested line fish as the small-harvested line fish were consistently bolder than the controls post the larval stages (8-22 dpf). Large- and small-harvested lines, but not control line, showed consistency in boldness through juvenile and subadult stages (46-85 dpf). Thus, both positive and negative size-selection fosters early emergence of collective personality with respect to the control line. All selection lines showed consistency in boldness in adult stages (105-190 dpf) and large-harvested line showed higher variability and plasticity in boldness compared to the other two lines. In presence of a live predator, the small-harvested line fish tend to be bolder and the large-harvested line fish where bolder at the highest risk of predation. Fish across all lines showed consistency in boldness, and the large-harvested line fish showed higher variability and lower plasticity in boldness compared to the other two lines.

We first found that boldness levels decreased significantly with increase in ontogenetic age across all selection lines. This meant that as fish matured from larval to the adult stages, they took significantly less risk to feed on the water surface following the unprecedented disturbance overhead due to sudden release and retrieval of the predator model. We expected this pattern only in the large harvested line where we predicted that fish would become shyer as adults but we found this across lines. The reason for decline in boldness with maturity is potentially based on the asset protection hypothesis according to which adults will be less prone to take risks for safeguarding possibilities for future mating while larval fish will take more risks since they have least investment in reproductive assets like gonads (Wolf et al. 2007). There could be other reasons behind these results. Firstly, the motivation for feeding increases in larval zebrafish after the absorption of yolk resulting in larvae to become voracious feeders. Adults with greater energy reserves and gut capacity may not require foraging as intensively as larval or juvenile fish (Fuiman and Webb 1988). Larval and juvenile fish have faster metabolic rates, lower body fat reserves and higher drag coefficients and therefore may be more inclined to take risks to feed than adults (Krause et al. 1998; Wootton 1994). Secondly, larval fish may perceive a looming stimulus like a predator model approaching overhead differently than adults (Fero et al. 2011). This could be possibly due to underdeveloped sensory and motor systems compared to adults. Larval fish may not show a significant startle response after the simulated threat and continue to swim on the surface unlike adults. Adults with developed visual and sensory systems are more vigilant, have a better knowledge of the risk-zone (surface) and may swim up to the surface only when hungry. Thirdly, adult zebrafish show more shoaling behaviour than larval or juvenile zebrafish (Fuiman and Webb 1988; Miller and Gerlai 2011). Since we tested fish in groups, larval and juvenile fish due to their low shoaling tendency had higher probabilities of leaving the association of groups for foraging. Contrarily, adults due to their increased shoaling tendency might have been more reluctant to leave the association of group for foraging. Our results contradict previous studies on mosquitofish that showed that juveniles showed decreased boldness compared to adults (Polverino et al. 2016a; Polverino et al. 2016b), and partially agree with a study on mangrove killifish (*Kryptolebias marmoratus*) where fish became bolder during early development followed by a reduction in boldness post sexual maturity (Edenbrow and Croft 2011). The reasons for this difference could be that these studies were conducted on individuals and did not measure boldness in the context of foraging. Also, species-specific differences and other methodological variations in the assay may be responsible.

We found that beyond the larval stages (8-22 dpf), the small-harvested line fish were significantly bolder than the control line fish while the large-harvested line fish did not differ in boldness compared to the controls. This supports our hypothesis that negative size-selection will lead to elevated boldness in zebrafish and that the increase in boldness will be more pronounced in the small-harvested line fish. The fact that the large-harvested line fish did not differ in boldness but the small-harvested line showed elevated boldness than the control line is in partial agreement with Andersen et al. (2018) which predicted that unless very large-sized fish are harvested, any kind of selective harvesting would foster boldness. Juveniles (46 dpf) of the small-harvested line were significantly bolder while the juveniles of the large-harvested line did not differ in boldness than the control line fish. These results are supported by previous findings of Uusi□Heikkilä et al. (2015) where juvenile individuals (30 day old) of the small-harvested line were bolder than juveniles of large-harvested and control lines though the study implemented an open-field assay to test risk-taking behaviour. Higher risk-taking to feed in juveniles in the small-harvested line could be justified by the need to develop energy reserves for investment in growth following the energy acquisition pathway (Enberg et al. 2012). On the other hand, no difference in boldness between large-harvested and control line fish would mean that though the large-harvested line evolved a fast life-history (Uusi□Heikkilä et al. 2015), this does not lead to higher foraging tendency to build energy reserves necessary for early gonadal investment. Further, we found that sub-adults (61-85 dpf) and adults (105-190 dpf) of the small-harvested line were significantly bolder while the large-harvested line did not differ in boldness compared to the controls. This agrees partially with our expectations that both lines would show elevated boldness. We saw that small-harvested line fish were bolder and that fish among all lines became shy as adults, as discussed before. The results are in agreement with a previous study by Sbragaglia et al. (2020) which implemented a similar assay to test group risk-taking among selection lines and found that boldness was higher among adults of the small-harvested line while the large-harvested line fish did not consistently differ in boldness compared to the control line fish. Our results are in contrast with the findings of Sbragaglia et al. (2019a) where individual adult females of the small-harvested line showed lower risk-taking tendency in an open-field test compared to the control line fish. An open field test measures exploratory behaviour rather than boldness (Réale et al. 2010) and the assay does not consider the vertical dimension of fish movement. Considering this dimension is important since zebrafish in holding are fed at the surface. The results with subadults and adults, like juveniles in the small-harvested line fish could be reasoned out based on energy acquisition mechanism (Enberg et al. 2012). Within the large-harvested line, we expected that fish would become shy as adults and our results support that. Though we see this pattern of change in boldness, this is not significantly different from the control line (Fig. 4a). Thus, our work does not support that size-selection alone leads to shyness. Life-history adaptations are stronger in the large-harvested than the small-harvested line fish (Uusi□Heikkilä et al. 2015) and therefore the large-harvested could be expected to differ significantly from the controls in behavioural traits. The fact that we do not see this despite a strong life-history evolution may be because of evolutionary resistance to change in behaviour. A study on medaka (*Oryzias latipes*) by Renneville et al. (2018) showed that life-history traits did not respond to anthropogenic (fishing like) selection at maturity even though the inverse anthropogenic selection had a strong impact on life history traits. Thus, selection may not always manifest changes in functional traits.

We did not find consistency in collective risk-taking to feed in larval stages (8-22 dpf) while we found significant behavioural consistency across juveniles (46 dpf) and sub-adults (61-85 dpf), and high consistency among adults (105-190 dpf). Consistency in boldness beyond juvenile stages (i.e. post 46 dpf age) was limited to the large- and small-harvested line fish which meant that size-selection fosters early emergence of collective personality. These results show that group risk-taking behaviour emerges as a collective personality trait in zebrafish only beyond the juvenile stage. Behavioural consistencies might change with increase in ontogenetic age because of development of motor and sensory capabilities (Fuiman and Webb 1988). Our results are similar to studies in eastern mosquitofish where personality differences emerged only in later stages (Polverino et al. 2016a; Polverino et al. 2016b), and compliment with studies in zebrafish that showed the development of shoaling behaviour with ontogenetic age (Buske and Gerlai 2011; Mahabir et al. 2013). In larval stages, fish do not rely on social information and have higher tendency to move away from groups and find resources on their own. This pattern changes with maturation when fish start shoaling resulting in more consistent behaviour. A study in killifish has shown that risk-perception is necessary for the development of personality (Edenbrow and Croft 2013), and reduced risk-perception among zebrafish from 8 to 22 dpf age could be another reason for the lack of behavioural consistency. Increase in behavioural consistency with ontogenetic age could be because of decrease in behavioural plasticity as predicted by theoretical models (Fawcett and Frankenhuis 2015; Fischer et al. 2014) and demonstrated in individual mosquitofish (Polverino et al. 2016a). We do not see this trend here probably because our study deals with groups, not individuals of zebrafish.

We see that behavioural variability increased from larval (8-22 dpf) to adult stages (105-190 dpf) among all selection lines. Behavioural plasticity increased for the small-harvested line, while increased and then decreased for large-harvested and control lines from larval (8-22 dpf) to adult stages (105-190 dpf). In the larval stages (8-22 dpf), the selection lines showed negligible among-group variability in boldness meaning that groups behaved similarly. We also found very low within-group variance among large- and small-harvested lines (but higher than control line) and this indicated low plasticity in behaviour. Therefore, larval fish groups across all lines generally took high risks to feed under simulated predation threat and were highly consistent in behaviour across time points. Our results on change in behavioural plasticity through ontogeny are in sharp contrast with previous studies on Atlantic molly *Poecilia formosa* (Bierbach et al. 2017) and eastern mosquitofish (Polverino et al. 2016a). These studies reported high behavioural plasticity among juvenile and low plasticity among adult individuals but we do not see this trend among groups of zebrafish. Through juvenile and subadult stages (46-85 dpf), the large-harvested line showed higher behavioural variability than small-harvested and control lines, and high behavioural plasticity. As adults (105-190 dpf), the large-harvested line showed higher behavioural variability and plasticity than small-harvested and control lines. These are in line with our expectations that positive size-selective mortality will foster higher behavioural variability and plasticity. This could be because of the internal conflict between reaping resources through foraging and avoiding being predated upon due to the smaller body-size. Our results align well with studies where fishery induced selection in pike (Edeline et al. 2009) and positive size-selective mortality in zebrafish (Uusi-Heikkilä et al. 2016) leads to higher trait variability. However, these studies dealt with morphological traits while our study demonstrates variability and plasticity in behaviour as a result of size-selection.

Under higher risks of predation, i.e. when zebrafish perceived visual and olfactory cues from a live fish predator, the small-harvested line showed higher risk-taking tendency (though not significantly) across all situations. This observation is in line with the results above and indicates that the small-harvested line always takes more risks to feed under any degree of predation threat. The large-harvested line fish were bolder at the highest predation risk i.e. when fish perceived both visual and olfactory cues from the cichlid fish. Thus, the large-harvested line fish demonstrated plastic adjustments of behaviour with respect to predation threat as expected. All selection lines showed significant behavioural repeatability, and the large-harvested line showed very high repeatability, indicating high consistency in behaviour across contexts and time points. This is similar to findings from previous studies where wild populations of zebrafish showed high consistency in risk-taking behaviour across contexts that differed in predation threat (Roy et al. 2017), and across time points (Roy and Bhat 2018), though the studies dealt with individuals. The large-harvested line showed higher behavioural variability and lower behavioural plasticity than the control line. This partly contrasts with previous results where we found lower behavioural plasticity in adults of large-harvested line. The high behavioural consistency in large-harvested line could be explained by low behavioural plasticity, as in eastern mosquitofish (Polverino et al. 2016a). Thus, positive size-selective mortality fosters higher variability in collective risk-taking similar to higher variation in body-size under different feeding environments seen previously (Uusi-Heikkilä et al. 2016), but low behavioural plasticity under high risks of predation. The small-harvested line showed lower behavioural variability and plasticity than the control line which means that negative size-selective mortality reduces variability and plasticity in boldness under high risks of predation. This partially agrees with previous results where adults of small-harvested line showed marginally higher behavioural variability than control line. Overall, the selection lines showed higher variability and plasticity in collective risk-taking in presence of live (aquatic) than simulated (aerial) predator.

## Conclusions

Our results demonstrated that five generations of intensive size-selective harvesting cause substantial changes in the emergence of collective personality. Collective personality emerges with increase in ontogenetic age among the selection lines of zebrafish supporting theoretical predictions (Fawcett and Frankenhuis 2015; Fischer et al. 2014) and empirical evidence from other species testing individuals (Edenbrow and Croft 2013; Polverino et al. 2016a). Personality is also stable within life-stages and across contexts as in other species (Cabrera et al. 2021). Negative size-selective mortality fosters increased boldness compared to other forms of selection under any kind of predation threat, be it aerial (simulated) or aquatic, but this difference is only evident beyond the larval stages. This elevated boldness can lead to increased natural mortality and be detrimental for populations, as predicted theoretically by Jørgensen and Holt (2013). Future research must experimentally test if this happens truly. Positive size-selection representing harvesting patterns commonly observed in most commercial and recreational fisheries does not foster change in collective risk-taking behaviour. However, positive size-selection fosters most plastic changes in boldness resulting in higher behavioural variability and varied levels of plasticity across different levels of predation risk. Despite these, a higher consistency in collective behaviour is observed in the large-harvested line. Negative size-selection results in decreased behavioural variability and plasticity than random size-selection. Our study not only provides empirical evidence for theoretical studies that predicted the direction of evolution of boldness with respect to size-selective harvesting (Andersen et al. 2018; Claireaux et al. 2018), but also shows whether and how selection patterns influence the ontogenetic development of boldness. Overall, size-selection alone favours boldness or no change in boldness, but does not promote shyness, in contrast to expectations of the timidity syndrome hypothesis (Arlinghaus et al. 2017). Thus, for shyness to evolve in response to fishing, selection must operate directly on behaviour. Future studies (like on learning, memory and cognition) are required to understand the adaptive significance of risk-taking behaviour among the selection lines.

## Acknowledgements

We sincerely thank David Lewis and Immanuel Kadlec for help with zebrafish breeding and maintenance, and Jelena Lewin for help with the experiments and scoring video data.

## Author’s contributions

TR and RA conceived the study; TR performed the experiments, analyzed the data, and wrote the manuscript with substantial inputs from RA.

## Ethics approval

This study was approved by State Office for Health and Social Affairs Berlin (LaGeSo), Germany (approval number: G 0036/21).

