## Supplementary Figure 1 for "Size-selective mortality fosters ontogenetic changes in collective risk-taking behaviour in zebrafish, *Danio rerio*"

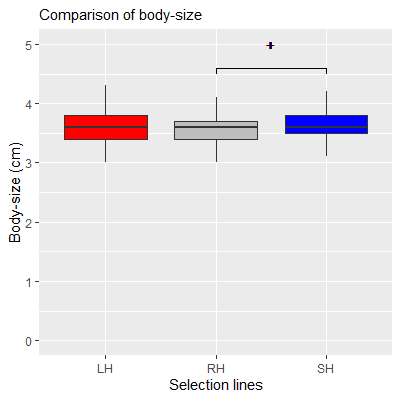

Figure 1: Comparison of body-size across selection lines in F_16_. The small-harvested line (SH) fish were larger than the control (RH) line fish. Significant differences is indicated with code **^+^** (p=0.08).
